# The highly evolvable nature of the antibiotic efflux protein TolC limits use of phages and bacterial toxins as antibacterial agents

**DOI:** 10.1101/2020.10.28.332486

**Authors:** Yusuf Talha Tamer, Ilona Gaszek, Marinelle Rodrigues, Fatma Sevde Coskun, Michael Farid, Andrew Y. Koh, William Russ, Erdal Toprak

## Abstract

Bacteriophages and bacterial toxins are promising antibacterial agents to treat infections caused by multidrug resistant (MDR) bacteria. In fact, bacteriophages have recently been successfully used to treat life-threatening infections caused by MDR bacteria [1–3]. One potential problem with using these antibacterial agents is the evolution of resistance against them in the long term. Here, we studied the fitness landscape of the *Escherichia coli* TolC protein, an outer membrane protein that is exploited by a pore forming toxin called colicin E1 and by TLS-phage [4, 5]. By systematically assessing the distribution of fitness effects (DFEs) of ~9,000 single amino acid replacements in TolC using either positive (antibiotics and bile salts) or negative (colicin E1 and TLS-phage) selection pressures, we quantified evolvability of the TolC. We demonstrated that the TolC is highly optimized for the efflux of antibiotics and bile salts. In contrast, under colicin E1 and TLS phage selection, TolC sequence is very sensitive to mutation. Our findings suggest that TolC is a highly evolvable target limiting the potential clinical use of bacteriophages and bacterial toxins.

## Main

TolC is an outer membrane protein conserved across Gram-negative bacteria and critical for the protection of bacterial cells against the toxicity of antimicrobial compounds such as antibiotics and bile salts (Figure 1A) [6–9]. TolC forms a homotrimeric channel, composed of a β-barrel domain that spans the outer membrane and a helical coiled-coil bundle that extends ~10 nm into the periplasmic space. TolC can partner with the AcrA-AcrB, MacA-MacB, and EmrA-EmrB protein pairs to form different efflux pump complexes in *E. coli* [10]. These efflux pumps render *E. coli* intrinsically resistant to several antibiotics such as β-lactams and macrolides [10, 11]. TolC also effluxes bile salts which are abundant in the human gut (Figure 1A) [12, 13]. Therefore, although *tolC* is typically not considered an essential gene for *E. coli*, the loss of *tolC* is costly in the presence of antibiotics or bile salts (Figure S1, S2). Interestingly, as TolC is exploited by both colicin E1 and the lytic TLS bacteriophage as a receptor, exposure to antibiotics (or bile salts) or colicin E1 (or TLS phage) creates opposing selective forces on maintaining the function of TolC, creating a convoluted fitness landscape where the evolutionary dynamics of TolC is highly unpredictable.

**Figure-1.**
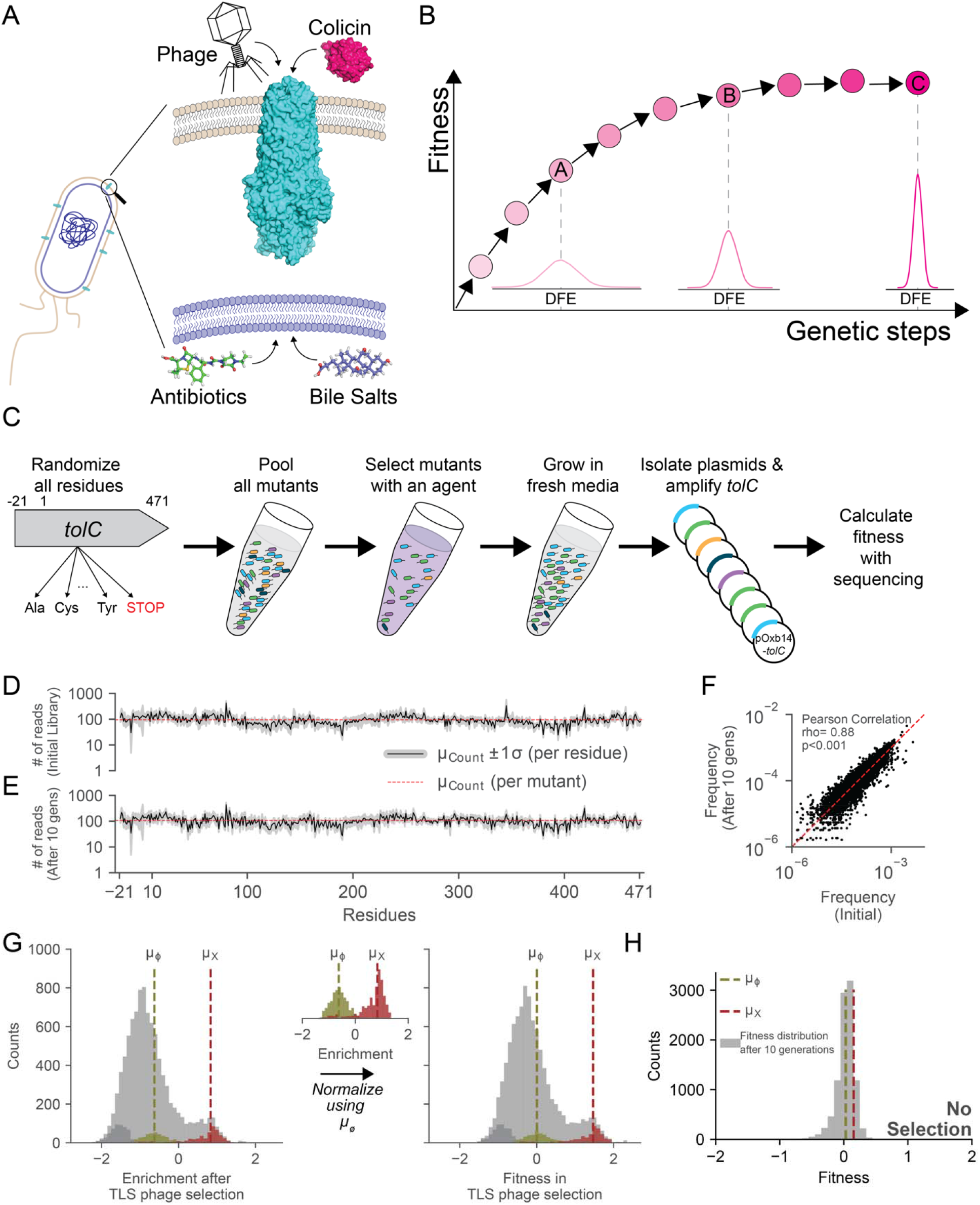
Creation of TolC mutant library. **(A)** TolC is an outer membrane protein in *E. coli* that is involved in the efflux of antibiotics and bile salts. TolC protein is also exploited as a receptor by TLS phage and colicin-E1, a bacterial toxin. **(B)** Fisher’s Geometric theorem predicts evolvability of biological systems using the widths of distributions of fitness effects (DFE). Large DFEs are indicators of high evolvability. Narrow DFEs are observed when biological systems are robust to genetic perturbations and the narrow width suggests lower potential to evolve. **(C)** Experimental procedure for whole gene saturation mutagenesis and fitness measurements. Deep Mutational Scanning of TolC was done by randomization of all 493 residues (including the signal sequence between residues −22 to −1, except the start codon) to 19 other amino acids and the stop codon. All mutants were pooled together and grown under selection of one of the four agents (antibiotics, bile salts, colicin-E1 or TLS phage). The *tolC* alleles were then harvested after a brief recovery growth in plain growth media, and frequency of mutations were calculated by deep sequencing of the *tolC* alleles. **(D-E)** Number of reads for all TolC mutations are plotted before (D) and after (E) growing the TolC mutant library for 10 generations (lower panel) in minimal media. Black line represents the mean (μ) number of reads per mutation at each residue. Gray area around the black line shows ±1 standard deviation from the mean. Horizontal dashed red line marks the mean number of reads for all TolC mutations. **(F)** Frequencies of all *tolC* mutations before and after 10 generations of growth in minimal media were highly correlated (Pearson, rho= 0.88 and p< 0.001). Red diagonal dashed line shows y=x line. **(G)** We calculated enrichment of each mutation by comparing mutation frequencies with and without selection (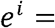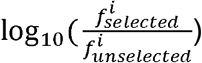; *e* stands for enrichment and *f* stands for frequency). Light green histogram represents enrichment values of mutations synonymous to the wild-type TolC protein sequence. Light red histogram represents mutants with early stop codons mutations. Mean values for the stop codon mutations (*μ*X) are represented with the vertical dashed red lines. Dark grey colored histograms on the left shows the mutants that went extinct after the selection. Their fitness values were calculated by assuming that their counts were equal to 0.01 after selection in order to manually separate them from the rest of the mutations. We used average fitness of synonymous wild type mutations (*μ*_∅_; dashed vertical green line) for defining relative fitness values of each mutation with respect to the wild type (WT) TolC sequence (*s^i^* = *e^i^* − < *e^WT^* >; s stands for fitness). **(H)** Distribution of fitness effects for the TolC mutant library after growth in minimal media (~10 generations) without any selection.

Mutational robustness is a common characteristic of evolved. For evolutionary success, a protein must tolerate spontaneous mutations for both survival and functional innovation. In the past decade, there have been many studies systematically assessing the fitness effects of mutations, particularly single amino acid replacements, on the function of proteins. Several of these studies have shown that protein function is robust to most single amino acid replacements [14–16]. In other cases, however, it was shown that many mutations can significantly deteriorate or even impair protein function [14–16]. As was originally proposed by Fisher, this problem gets even more complex because of the pleiotropic effects of mutations that can improve or worsen multiple traits simultaneously [17]. To date, most studies of the evolutionary dynamics of proteins have focused on the selection pressure imposed by a single, specific growth condition, although the natural process often involves multiple, potentially opposing selection pressures [18–20]. This difference limits our understanding of how functional protein sequences have adapted to environmental fluctuations.

According to Fisher’s fundamental theorem, the distribution of fitness effects (DFE) for mutations can be used as a metric for protein evolvability [17]. Using this theorem, we have previously shown that the width of the DFE is a good predictor for the rate of evolution of antibiotic resistance [21]. If a biological system is highly fit and robust to genetic perturbations, mutations are expected to have small fitness effects yielding a narrow DFE, centered around neutrality (Figure 1B, point C). However, if the biological system has low fitness and is sensitive to genetic perturbations, mutations are expected to have larger fitness effects and hence the DFE will be wider (Figure 1B, points A and B). Of note, in Figure 1B, we use normal distributions with varying widths to represent the DFEs, but realistically there is no way of predicting the shapes of these distributions and there may be outlier groups of beneficial or deleterious mutations.

In this study, we explore the evolvability of the efflux protein TolC using a saturation mutagenesis library which contains all possible amino acid replacements for each position of the TolC protein (barring the start codon). Utilizing both positive and negative selection, we performed a deep-sequencing based fitness assay to quantify the fitness landscape of TolC. We **systematically assesed the distribution of fitness effects (DFEs) of ~9,000 single amino acid replacements in TolC under antibiotics, bile salts, colicin E1, or TLS-phage selection. We demonstrated that TolC is highly optimized for the efflux of antibiotics and bile salts. In contrast, under colicin E1 and TLS phage selection, TolC sequence is very sensitive to mutations.** This observation is important in the context of public health where agents such as bacteriophage and bacterial toxins are favorably viewed as less mutagenic alternatives to antibiotic therapy.

## Results

### Creation of a TolC mutant library

We measured the evolvability of TolC by quantifying DFEs of all possible single amino acid replacements in the presence of four physiological stress factors: antibiotics (piperacillin-tazobactam), bile salts, colicin E1, and TLS phage (Figure 1C-H). First, we generated a *tolC* deletion strain (*E. coli*-Δ*tolC*, Methods) which became more sensitive to both antibiotics and bile salts (Figures S1 and S2) relative to its wild-type parent strain (BW25113) [22, 23]. The *tolC* deletion strain was also more resistant to both colicin E1 and TLS phage relative to its wild-type parent (Figure S1). We reintroduced the *tolC* gene into this strain using a plasmid that has a constitutively active promoter (pSF-OXB14, Oxford Genetics) and rescued both the antibiotic and bile salt resistance and the colicin E1 and TLS phage sensitivity of the *E. coli*-Δ*tolC* strain (Figure S1 and Figure S2). We mutated all residues except the start codon (471 residues in the mature TolC protein, and the 21 residue-long signal peptide) of TolC and generated a pool of ~9,841 (492 sites x 20 aa and a stop codon) mutants (Figure 1C, Figure S3). We cloned the mutated *tolC* genes into the pSF-OXB14 plasmid and then transformed the *E. coli*-Δ*tolC* strain with this pool of plasmids carrying mutated *tolC* genes. We randomly selected 30 amino acid positions from our library and using Sanger sequencing, confirmed that all 30 of the mutated sites were randomized and these *tolC* alleles did not have unintended mutations at other sites (Figure S3). For amplicon sequencing, we pooled mutants into five sub-libraries and carried out parallel selection and sequencing experiments (Methods). We deep-sequenced the *tolC* genes in each sub-library by utilizing the Illumina MiSeq platform and verified that 98.9% of possible amino acid replacements in the mutant library yielded at least 10 counts when sequenced (Figure 1D-E), with ~1,800 reads per residue or an average of ~90 reads per amino acid replacement (Figure 1D-E). We also confirmed that frequencies of the mutations in the *tolC* library did not change significantly when the library was grown in growth media without selection (Figure 1F, ρ=0.88, p<0.001, Pearson correlation).

### Systematic quantification for fitness effects of TolC mutations under positive or negative selection

We measured fitness effects of TolC mutations under selection using a liquid based assay (Figure 1C). In brief, we grew mutant libraries in growth media to saturation, diluted them to an OD600 of 0.001, and then grew these cultures in the presence of one of the four selection factors for three hours. Cells were then washed and grown in nonselective media for six hours. Finally, we harvested plasmids carrying *tolC* mutants and performed amplicon sequencing to count the surviving *tolC* variants(Methods). The duration of selection and recovery periods were optimized to maximize the dynamic range of the measurements and to minimize the chances of losing some alleles during plasmid harvesting (Methods, Figure S1 and S4). Of note, all concentrations used in these assays were above the minimum concentrations sufficient to kill wild-type *E. coli*, except the bile salts. The maximum soluble amount (50 mg/ml) of bile salts in our selection experiments inhibited growth of the *E. coli*-Δ*tolC* strain but not the wild-type *E. coli* strain (Figure S2). A control experiment with no selection was performed in parallel, to decouple fitness effects due to growth defects.

For calculating fitness, we determined the enrichment of each mutation by comparing mutation frequencies with and without selection (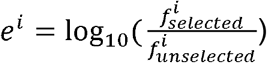, where *e* represents the enrichment, and *f* the frequency, of mutation *i*, Figure 1G). We used the average fitness of synonymous wild-type mutations as a reference point for defining relative fitness values (*s*) of each mutation with respect to the wildtype (WT) TolC sequence (*s^i^* = *e^i^* − < *e^WT^* >; Figure 1G, green bins). As a control, we compared the fitness effects of early stop codons (Figure 1G, pink bins) with the *E. coli*-Δ*tolC* strain supplemented with the *tolC* gene (Figure 1G, green bins) and confirmed that the results we obtained using our sequencing-based assay matched our observations in batch culture (Figure S4) both qualitatively and quantitatively. By comparing the enrichments of mutations in the absence of selection (Methods) relative to the frequencies of mutations in the library before any growth or selection, we confirmed that TolC mutations did not have significant fitness effects in the absence of selection (Figure 1F, H).

Figure 2A-B shows the fitness effects for a subset of single amino acid replacements in TolC in the presence of antibiotics (6 μg/ml), and TLS phage (2.5 x 10^8^ pfu/ml). Figure S5 summarizes the fitness effects of the entire mutation library under all four selection factors. We found that the fitness effects of mutations in the presence of antibiotics or bile-salts were mostly neutral (Figure 2A and Figure S5A-B, white pixels) except a group of mutations increasing sensitivity to antibiotics or bile salts (Figure 2A blue pixels, Figure 2 C-D insets). Figure 2C and Figure S6 shows the corresponding DFEs. When we repeated the same assay using 10 times lower dose of antibiotics (0.6 μg/ml, which is still higher than the MIC value of piperacillin-tazobactam for wild-type *E. coli*, Figure S2A) and bile salts (5 mg/ml), we saw that the DFE in bile salt selection did not change much but the DFE in antibiotics became slightly narrower further verifying the robustness of the TolC sequence under antibiotic selection (Figures 2D and S6). None of the mutations increased resistance to either antibiotics or bile salts suggesting that the *E. coli* TolC sequence is highly optimized for the efflux function as the TolC sequence is mostly insensitive to mutations under these conditions, evident from the corresponding DFEs (Figure 2C, D and Figure S6A, B).

**Figure-2.**
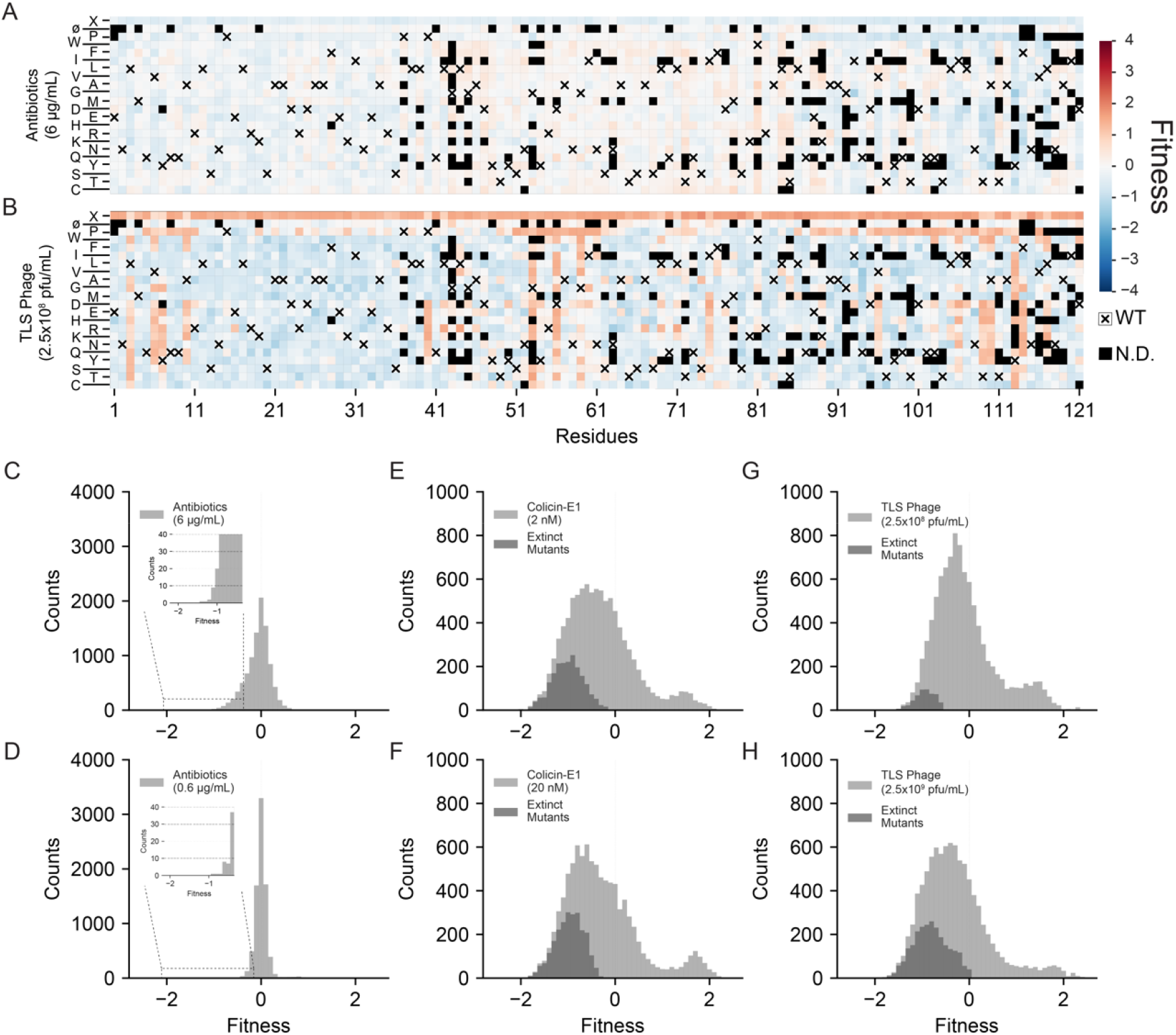
Mutant library selection using antibiotics and TLS Phage. Heatmaps summarizing a subset of fitness values of TolC mutations under selection of (A) Antibiotics (6 μg/mL), (B) TLS Phage (2.5×10^8^ pfu/mL). Columns within each matrix represent the TolC residues (from residue 1 to 121), and columns represent mutations to other codons causing synonymous(∅), nonsynonymous, or nonsense (X) mutations. Protein sequence of the wild-type TolC are represented by white pixels with cross on them. Mutants with low read counts (threshold= μ-1.5⍰) in the initial TolC library or after growth in plain media (untreated) were excluded from fitness calculations and represented by black pixels. All other calculated fitness values were colored from dark blue (increased susceptibility) to white (neutral effects) to dark red (increased resistance). Top row of each heatmap shows the effect of early stop codon mutations (X). Second rows from the top show the effect of silent mutations synonymous to native codon in TolC. Under antibiotics selection (A), nonsense mutations increased susceptibility, whereas under TLS phage selection (B), nonsense mutations increase resistance. **(C-H)** Distribution of fitness effects (DFEs) for different selection agents. For every selection agent, DFEs were calculated under two different selection strengths. DFEs under antibiotic selection are narrow and centered around neutrality (s = 0) regardless of the selection strength, with tails extending to the left (increased sensitivity, insets). Under both colicin-E1 and TLS phage selections, DFEs were wide and mean fitness effects of mutations were negative, suggesting that TolC was not robust to mutations under selection to these agents. Under both colicin-E1 and phage selection, many mutations were initially present in the TolC library but went extinct after the selection (dark gray bins). Although it is not possible to calculate fitness in these cases, in order to show them on the histograms, we set their final counts to 1 (pseudocount) and calculated a fitness value such that they are still visible. Under both colicin-E1 and phage selections, DFEs were bimodal with a second peak corresponding to mutations increasing resistance. Means and standard deviations of light gray histograms are tabulated in Table S1.

On the contrary, in the presence of colicin E1 or TLS phage selection, the majority of the TolC mutations had large effects on bacterial fitness (Figure 2B, Figure S5C-D, blue pixels) sometimes making *E. coli* cells more susceptible to colicin E1 or phage induced death. A subset of mutations increased bacterial resistance to colicin E1 or TLS phage (Figure 2B, red pixels). Corresponding DFEs were wider relative to the DFEs in antibiotics and bile salts (Figure 2E-H, Table S1). We repeated these measurements using ten times higher concentrations of colicin E1 and TLS phage and showed that the DFEs under these conditions were still wide (Figure 2F,H). These observations suggested that, under colicin E1 or TLS phage selection (Figure 2 E-H), the *tolC* gene has the potential to evolve resistance and resides at a sub-optimal fitness state as the TolC sequence is very sensitive to mutations.

### Relationship between strength of selection and fitness effects

We measured fitness effects of TolC mutations using different doses of colicin E1 in order to measure the relationship between mutational sensitivity and selection strength. In these experiments, we used increasing concentrations of colicin E1 (0, 5pM, 0.1 nM, and 2 nM, Figures 3A-E). In addition, we measured fitness effects of TolC mutations in the presence of TLS phage particles (Figure 3F) and bile salts (Figure S6C). These measurements were done using the Illumina NovaSeq platform and yielded nearly hundred-fold higher number of reads compared to the MiSeq platform. As the NovaSeq platform provided large number of sequencing reads, we did not observe any extinct mutations and we were able to quantify fitness values with greater confidence. We found that, as the selection strength by colicin E1 increases, the mean values of the DFEs shift to more negative values and the widths of DFEs become larger (standard deviation, Figure 3A-E). Similarly, the DFE under phage selection was still wide (Figure 3F) despite the use of 10-fold fewer phage particles compared to our previous measurements (Figure 2), in agreement with our observations using the MiSeq platform. On the contrary, the DFEs under bile salt (5mg/ml) selection was narrow, similar to the DFEs under no selection (Figure S6). Finally under both colicin E1 and phage selection, we found that a considerable fraction of TolC mutations were resistance-conferring mutations (Figure 3A-D, F-G, magenta). Almost half of these mutations were early nonsense substitutions (46% for both colicin E1 selection and TLS phage selection) that also induced antibiotic or bile salt sensitivity due to disruption of the efflux machinery. When we excluded stop codon mutations, there were still many (372 mutations spanning 168 residues for colicin E1 selection, 408 mutations spanning 184 residues for TLS phage selection) resistance-conferring mutations suggesting that the TolC sequence was only one mutation away from developing resistance to colicin E1 or TLS phages (Figure 3G). Using phages or colicin E1 in combination with antibiotics may potentially reduce the rate of evolution to some extent as early stop codon mutations will be eliminated by the use of antibiotics. However, many resistance conferring mutations will still be available and extended use of phages or colicin E1 in clinical settings will lead to selection of resistant TolC mutants, limiting the success of these therapies in the long-term.

**Figure 3.**
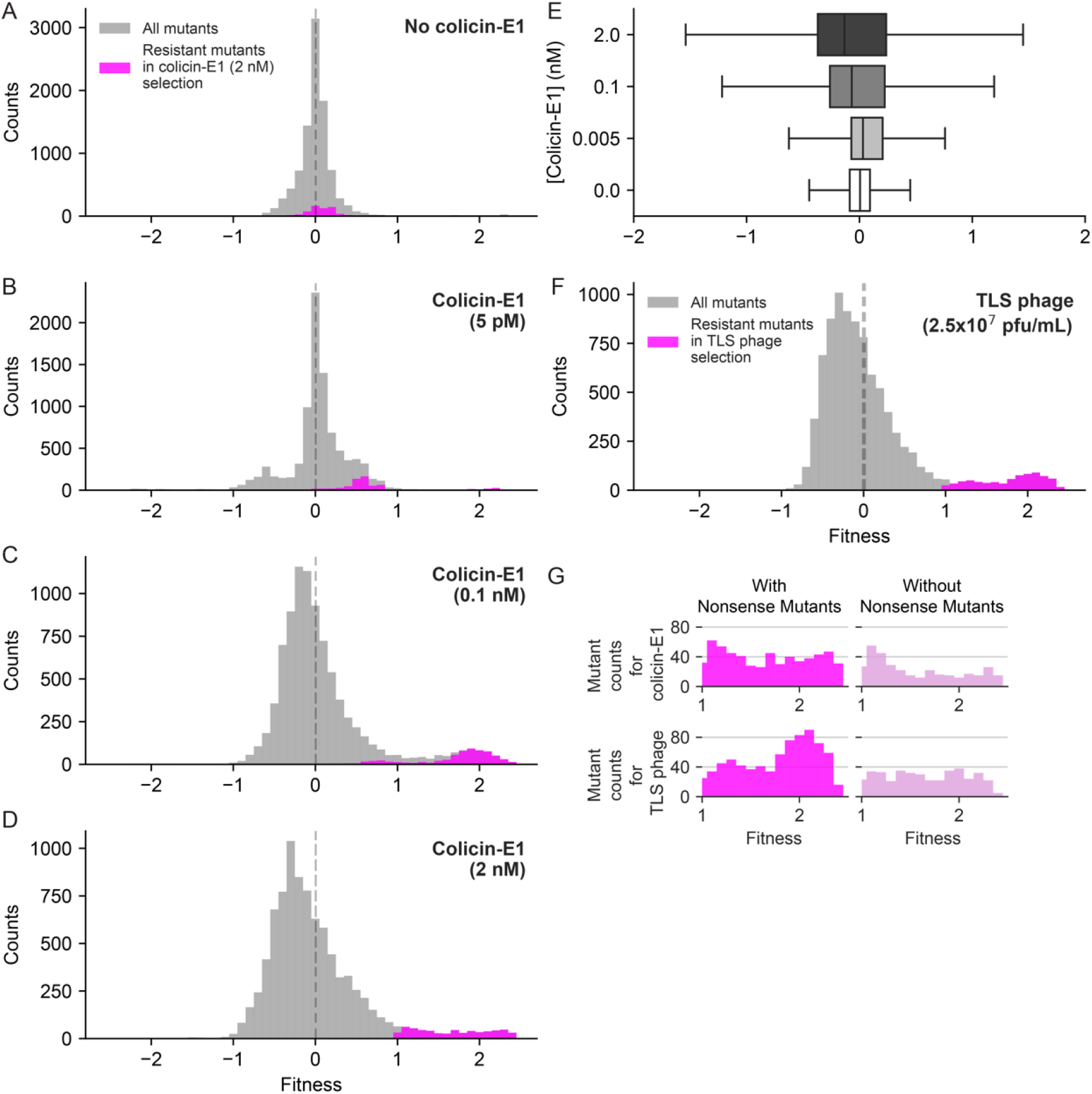
Impact of selection strength on DFE. **(A-D)** Selection strength alters the shape and **(E)** width of the distribution of fitness effects under colicin-E1 selection. We measured fitness effects of TolC mutations by Illumina NovaSeq sequencing platform which yielded ~100 times more reads per mutation, compared to Illumina MiSeq platform, and increased resolution of our fitness measurements. DFEs for TolC mutations under selection with **(A-D)** increasing concentrations of colicin-E1, and **(F)** TLS Phage (2.5×10^7^ PFU/mL). Magenta colored bins in panels A to D highlight resistance-conferring mutations that had fitness values larger than 1 (10-fold change in frequency) under selection with 2nM of colicin-E1. Magenta colored bins in panel F highlight resistance-conferring mutations that had fitness values larger than 1 under phage selection. **(G)**(left) Histograms of all resistance-conferring mutations under colicin-E1 (2nM, 685 mutations) selection and TLS phage selection (761 mutations). (right) Histograms of all resistance-conferring mutations, excluding stop codon mutations, under colicin-E1 (2nM, 372 mutations) selection and TLS phage selection (408 mutations).

The average fitness effects of TolC mutations under bile salt and antibiotic selection were both very small and weakly correlated (Figure S6D, ρ= 0.28 and p<0.001 Pearson Correlation), making it difficult for us to determine whether efflux efficiencies of these molecules by TolC were controlled by similar mechanisms. However, the fitness effects of TolC mutations under phage and colicin E1 selection were broad and significantly correlated (Figure-4A, ρ= 0.62 and p<0.001, Pearson Correlation) suggesting shared infection mechanisms by these agents. As expected, fitness effects under selection by neither TLS nor colicin E1 showed significant correlation with those measured in the presence of antibiotics (Figure 4B,C).

**Figure 4.**
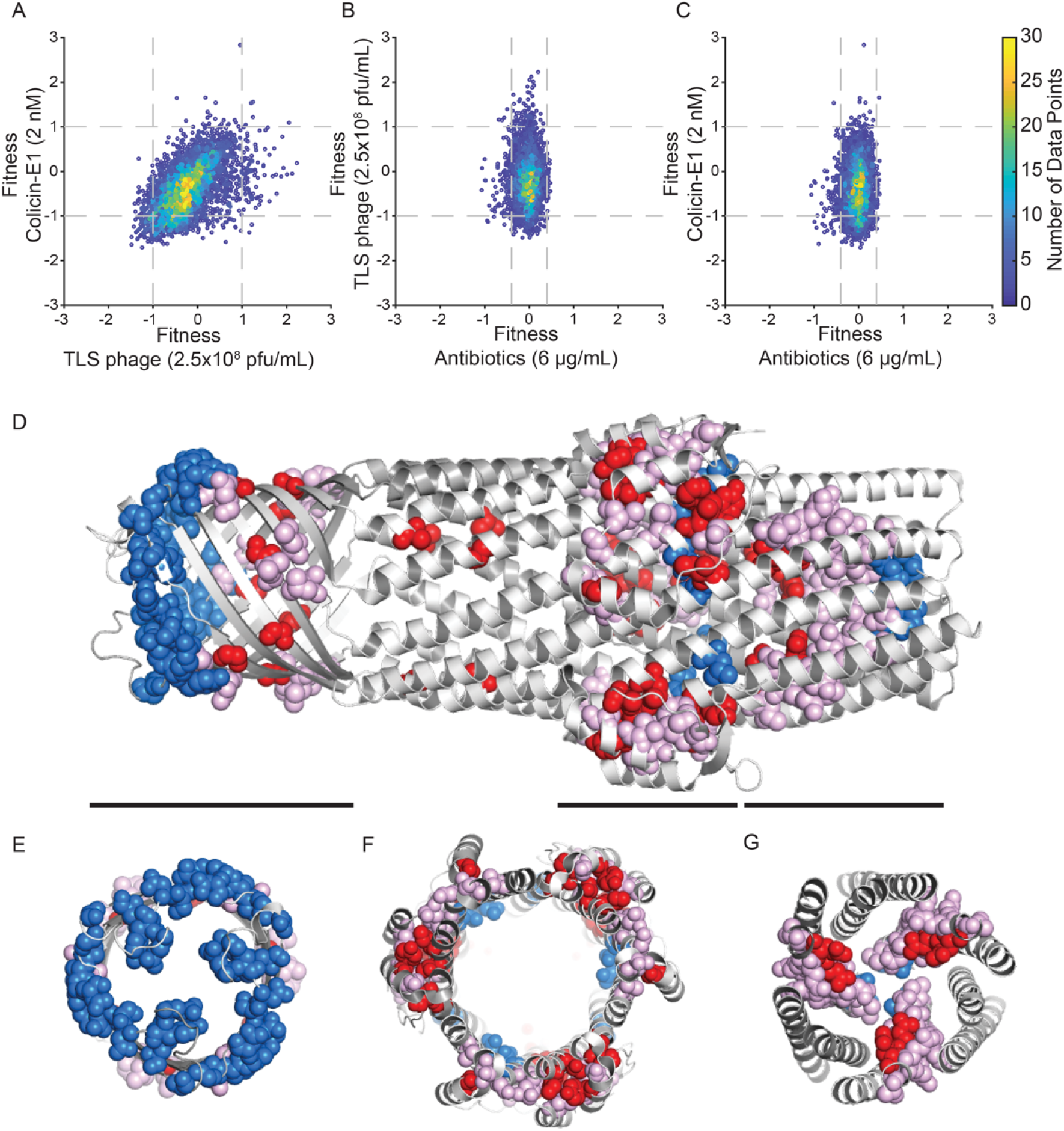
Analysis of mutations from selection experiments. **(A-C)** Comparison of fitness effects of TolC mutations under different selections. These fitness values correspond to consistent values used to plot histograms in Figure 2C-H. Horizontal and vertical dashed lines indicate the significance thresholds (± 1.5 standard deviations) for each selection condition, which correspond to ~10 fold increase or decrease in frequency relative to wild-type TolC (fitness values >+1 and <−1) under colicin-E1 or TLS phage selection. Significance threshold under antibiotic selection were ~2.5 fold increase or decrease in frequency relative to wild-type TolC (fitness values > +0.4 and <−0.4). **(D)** Side-view of the TolC trimer. The 47 most sensitive residues (representing the top 10%) for TLS and colicin E1 are shown in spheres. Positions that show sensitivity to TLS only are colored blue, sensitivity to colicin E1 only in red, and sensitivity to both perturbations in pink. Black bars correspond to slices along the pore axis, shown in **(E-G)** highlighting the three structural regions where sensitive residues are located.

### Structural mapping of mutation-sensitive TolC residues

Comparison of the mean fitness effects of TolC missense mutations in the presence of colicin E1 and TLS phage revealed that resistance to these factors can arise from mutations in three distinct structural regions (Figure 4D-G), comprising the extracellular surface of the beta-barrel domain, residues near the so-called “equatorial domain”, and residues in the periplasmic pore opening [24]. While positional sensitivity to TLS and colicin E1 selection showed strong overlap, there were several residues that were more sensitive to one of the two perturbations. Most notably, mutations in a cap of residues on the extracellular surface of the beta-barrel domain caused resistance to TLS phage, but were relatively insensitive to colicin E1 (Figure 4D, E). Residues near the equatorial domain and in the periplasmic pore exhibited sensitivity under both conditions (Figure 4F, G). The effects of mutations under different selection factors may provide some insight into the mechanisms by which phage and colicin E1 exploit TolC for entry into the cell.

Looking at individual mutations, rather than the mean effects averaged over each position, provided additional insight. We considered mutations that had fitness effects larger than three standard deviations from the mean (Methods) and excluded early stop codon mutations that were equivalent to loss of the *tolC* gene. For this analysis, we also excluded mutations that did not have consistent fitness effects between experimental replicates (Methods). We grouped TolC mutations that passed these criteria as summarized and highlighted mutated residues on the TolC monomer (Figure S7). Interestingly, many of these residues clustered together suggesting that they might induce similar changes on the TolC structure due to their physical proximity. We found a cluster of residues that made *E. coli* cells resistant (Figure S7, dark red residues) to both colicin E1 and TLS phage when mutated, without significantly changing antibiotic or bile salt resistance. Strikingly, there was an independent cluster of mutations, that made *E. coli* cells more sensitive (Figure S7, dark blue residues) to both colicin E1 and TLS, without significantly changing antibiotic or bile salt resistance. One plausible explanation is that mutations in the helical coiled-coil bundle extending into the periplasmic space (Figure S7, highlighted in dark red and dark blue) are altering the interactions of TolC with other proteins involved in colicin E1 or phage entry. Although not much is known about TLS entry mechanism into bacterial cells, at least three other proteins are reported to be involved in colicin E1 entry and translocation (TolA, TolQ, and TolR). Mutations in the residues on the beta barrel (Figure S7, highlighted in dark red) might be increasing resistance by weakening colicin E1 and TLS binding without significantly altering antimicrobial efflux. Mutations in the residues highlighted in cyan (Figure S7) show increased resistance to colicin E1 and TLS phage while making bacteria more sensitive to both antibiotics and bile salts. This effect can be explained by misfolding of TolC or a possible constriction of the TolC channel that mechanically or electrostatically obstructs the passage of all antibiotics, bile salts, colicin E1, and TLS phage (or viral DNA).

## Discussion

Utilization of bacterial toxins and bacteriophages to fight bacterial infection has been proposed for decades and was successfully used in some life-threatening infections. In fact, bacteriophages that bind the *Pseudomonas aeruginosa* OprM, an efflux protein homologous to TolC, were recently used to treat a patient with a life threatening pan-resistant *P. aeruginosa* infection [2]. Our results suggest that if evolutionary aspects are not taken into account, treatments using phages and bacterial toxins are prone to failure in the long term, consistent with rapid evolution of phage resistance during treatment of this patient (Personal communication with Ryland Young, Texas A&M University). Our study also provides structural clues for understanding mechanisms of TolC mediated efflux, colicin E1 binding and translocation, and bacterial infection with TLS. Future structural and functional studies investigating interactions of TolC with its partners and the spatiotemporal dynamics of these interactions will help to define strategies for controlling evolutionary outcomes, a key step in addressing problems such as antibiotic and drug resistance.

## Acknowledgements

We thank Dr. Michael Stiffler for the helpful discussions on making the saturation mutagenesis library. We also thank Dr. William A. Cramer, and Dr. Joe Fralick for providing Colicin E1 plasmid and TLS phage. We thank Dr. Richard Neher and Xiaowei Zhan for their advice on sequence analysis. We thank Ayesha Ahmed for colicin E1 purification and help with sequencing experiments.

## Online Methods

### Growth Media and Strains

*E. coli* cells were grown at 37°C in M9 minimal medium (248510, Difco) supplemented with 0.4% glucose (50-99-7, Fisher Scientific) and 0.2% amicase (82514, Sigma). BW25113 wild-type *E. coli* strain (CGSC#: 7636) and the Δ*tolC732::kan E. coli* strain (CGSC#: 11430) were obtained from the Coli Genetic Stock Center. Kanamycin resistance marker was removed from the Δ*tolC732::kan E. coli* strain following the protocol in reference [22]. This strain is referred as the Δ*tolC* strain throughout the manuscript. We whole-genome sequenced both the wild-type (BW25113) and the Δ*tolC E. coli* strains and confirmed that no other mutations besides the *tolC* deletion were present in the Δ*tolC* strain.

### Saturation mutagenesis assay for the *tolC* gene

pSF-Oxb14 plasmid was obtained from Oxford Genetics (OGS557, Sigma). This plasmid contained a kanamycin resistance cassette and an Oxb14 constitutively open promoter region. The *tolC* gene was PCR amplified from the BW25113 (wild-type) strain using 5’<u>ATTCAAAGGAGGTACCCACC</u>ATGAAGAAATTGCTCCCCATTC-3’ (forward), and 5’<u>AGAAATCGATTGTATCAGTCTCAG</u>TTACGGAAAGGGTTATGAC-’ (reverse) primers. It was then cloned into the pSF-Oxb14 plasmid using the NEBuilder HiFi DNA Assembly Kit (E5520, New England Biolabs), following the protocol provided by the manufacturer. Bold and underlined nucleotides in primer sequences overlap with the plasmid sequence. The integrated *tolC* gene was confirmed to have no mutations by Sanger sequencing.

Whole gene saturation mutagenesis was performed by two PCR reactions individually for each codon in the *tolC* gene, including the first 22 amino acid long signal sequence. First PCR reaction amplified a portion of the *tolC* gene in the pSF-Oxb14-*tolC* plasmid and randomized the targeted codon with a primer that contained a randomized NNS nucleotide sequence (N stands for A, C, G, or T nucleotides and S stands for G or C nucleotides) for the targeted codon (this PCR product is referred as insert). Second PCR reaction amplified the rest of the pSF-Oxb14-*tolC* plasmid (this PCR product is referred as backbone). Our custom software for designing mutagenesis primers is available at https://github.com/ytalhatamer/DMS_PrimerDesignTool. Inserts were cloned onto the backbones using the NEBuilder HiFi DNA Assembly Kit (E5520, New England Biolabs and assembled plasmids were transformed into NEB-5-alpha (C2987, New England Biolabs) cells. Plasmid extraction from these cells was done using Nucleospin Plasmid kit (740588, Macharey-Nagel). As this assay produced libraries per each residue, plasmid concentrations were measured and then equimolar amounts of each library were pooled into five sublibraries for 2×250bp paired-end MiSeq sequencing (residues 2-110, 90-210, 190-310, 290-410, 390-493) and twelve sublibraries for 2×150bp paired-end NovaSeq sequencing (1-40, 36-82, 79-124,120-166, 162-208, 204-250, 246-292, 288-334, 330-376, 372-418, 414-460, 456-493). Finally, these pooled sublibraries were transformed into Δ*tolC* strain for selection experiments. All growth and selection assays with the library were done using 50 μg/ml kanamycin in minimal M9 media.

### Colicin E1 purification

A colicin E1 expression vector with IPTG inducible T7 polymerase promoter was kindly provided by Dr. William A. Cramer (Purdue University). Only Colicin E1 was amplified and put back to an empty pET24a plasmid to remove immunity protein. Plasmids were then transformed into BL21-DE3 cells for expression and purification. Cells were grown in TB broth media and colicin E1 was purified first with a size exclusion chromatography. Elutes corresponding to the size of Colicin E1 (~57 kDa) were further purified using a cation exchange chromatography in Sodium borate buffer with a salt gradient of 0-0.3M (NaCl). All fractions are collected and analyzed by SDS-PAGE. Elutes with right band sizes pooled and concentrated using Amicon Centrifugal filters with 30K pore size (UFC803024, Milipore).

### TLS Phage Harvesting

TLS phage strain was kindly provided by Dr. Joe Fralick (Texas Tech University). Phage propagation and purification were done following the protocol described in [25]. Briefly, overnight grown bacterial cells were diluted hundred times in 100 mL of LB medium with 5 mM CaCl_2_ and incubated 2-3 hours till optical density reached 0.4-0.6. Phage particles were added to the culture and the culture was shaken (at 37°C) until the culture became optically clear. Cell lysates were spun down in 50 mL falcon tubes at 4000 x g and for 20 minutes. Supernatant was filter sterilized using 0.22 μm filters. Chloroform was added to the filtered phage solution (10% v/v final chloroform concentration) and the solution was vortexed shortly and incubated at room temperature for 10 minutes. Finally, the phage lysate and chloroform mixture were centrifuged at 4000 x g for 5 minutes. Supernatant was removed, aliquoted, and stored at 4°C.

### Selection Assay

We used Piperacillin-Tazobactam (NDC 60505-0688-4, Apotex Corp), bile salts (B8756, Sigma-Aldrich), colicin E1, and TLS phage in selection assays. TolC mutant sub libraries were separately grown overnight in M9 minimal media supplemented with 50 μg/mL kanamycin. These cultures were diluted to the optical density of 0.001 in 10 mL of M9 minimal media supplemented with 50 μg/mL kanamycin (~5×10^6^ cells). Selection agents were added to each sub library and cultures were incubated at 37°C for three hours. All cultures were spun down at 7000 x g for 2 minutes and pellets were resuspended in fresh M9 minimal media supplemented with 50 μg/mL kanamycin. These cultures were then incubated at 37°C with shaking for six hours. Following this step, libraries were centrifuged at 7000 x g for 2 minutes and pellets were collected for plasmid purification. Different regions of the *tolC* genes were amplified with PCR and indexed using Illumina Index sequences (Supplementary Data). These regions were spanning residues 2-110, 90-210, 190-310, 290-410, and 390-493 for 2×250bp paired-end MiSeq sequencing. For 2×150 bp paired-end NovaSeq sequencing, we amplified and indexed the residues 2-40, 36-82, 79-124,120-166, 162-208, 204-250, 246-292, 288-334, 330-376, 372-418, 414-460, and 456-493.

### Sequence analysis

Paired ended sequencing reads were first merged using the FLASh tool [26] (Customized parameters: -m 40 -M 100). Reads covering primers overlapping with the upstream and downstream of the amplified regions of *tolC* were excluded. Sequence reads were compared to the wild-type *tolC* sequence and mutations were listed. Sequence reads that had mutations in more than one residue were excluded from the analysis. Synonymous mutations yielding the same amino acid replacement were grouped together. Frequency of each mutation was calculated by dividing number of counts of for that mutation with number of all reads, including alleles with multiple mutations 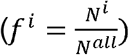. For calculating fitness, we first determined enrichment of each mutation by comparing mutation frequencies with and without selection 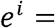 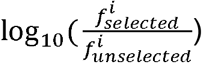; e stands for enrichment and f stands for frequency, Figure 1G). Since randomized mutations of each residue created traceable mutations synonymous to the wild-type protein sequence, we were able to use average fitness of synonymous wild-type mutations for defining relative fitness values of each mutation with respect to the wild-type (WT) TolC sequence (*s^i^* = *e^i^* − < *e^WT^* >; s stands for fitness, Figure 1K, green bins). As a sanity check, we compared the fitness effects of early stop codons with the phenotype of the *E. coli*-Δ*tolC* strain (Figure 1G, pink bins) and confirmed that the results we obtained our sequencing-based assay matched our observations in batch culture (Figure S4) both qualitatively and quantitatively. By comparing the enrichments of mutations in the absence of selection relative to the frequencies of mutations in the library before any growth or selection, we confirmed that TolC mutations did not have significant fitness effects in the absence of selection (Figure 1F, H).Our source code for data analysis is available at https://github.com/ytalhatamer/DMS_DataAnalysis.

## Supplementary Figures

**Figure S1.**
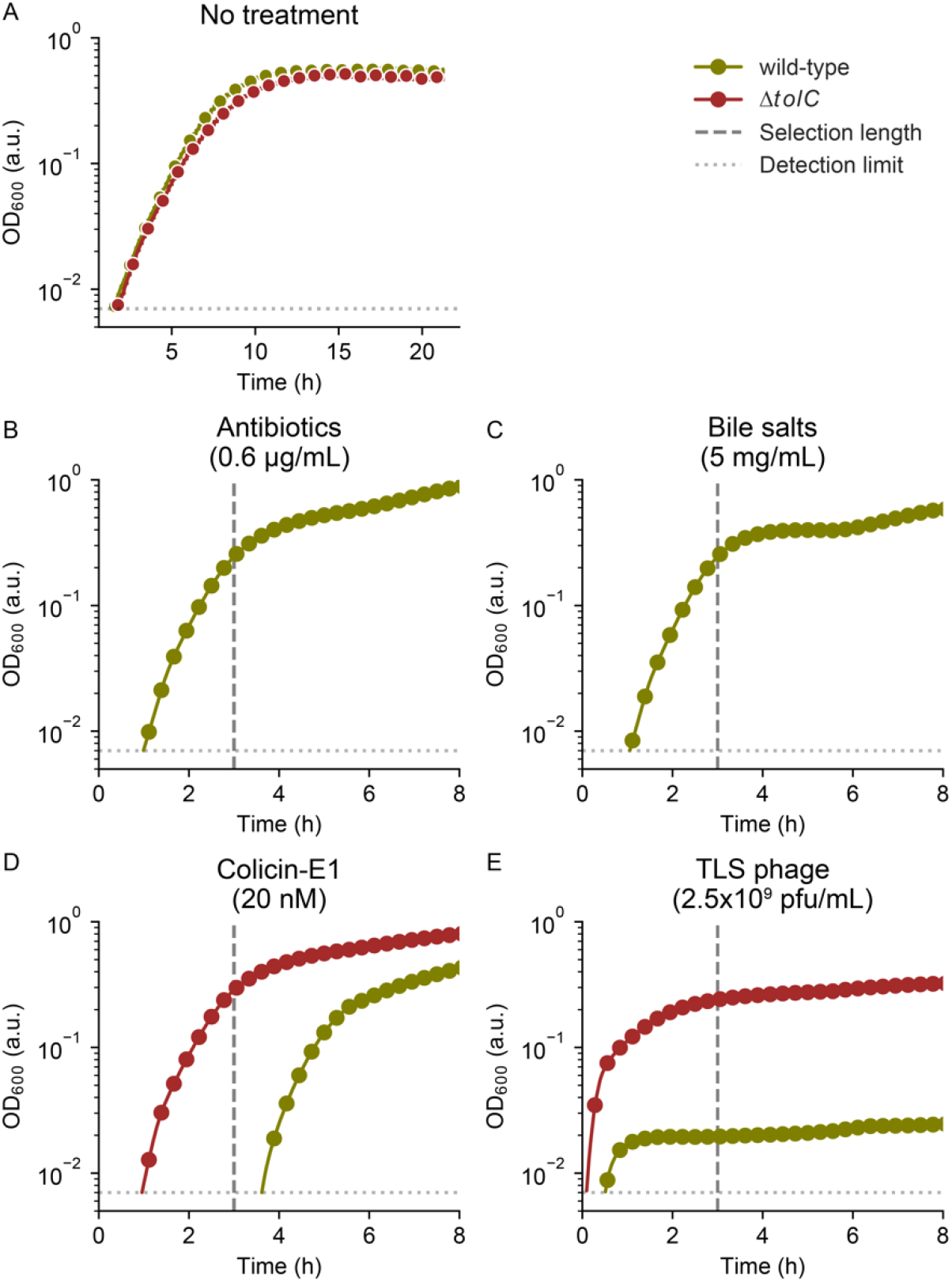
Deletion of the *tolC* gene is costly under antibiotic and bile salt selection and beneficial under colicin-E1 and TLS Phage selection. **(A)** Growth curves of wild-type and ΔtolC strains without selection pressure. Dark green colored lines represent growth curves of the wild type (BW25113) *E. coli* strain. Dark red colored lines represent growth curves of the BW25113 E. coli strain with *tolC* gene deletion (Δ*tolC*). In our fitness assays, we used a duration of three hours for selection (vertical grey dashed line) in order to maximize the fitness difference between the wild type *E. coli* and *E. coli*:Δ*tolC*. Horizontal gray dotted line represents the detection limit of the spectrophotometer used (OD_600_: 0.007).

**Figure S2.**
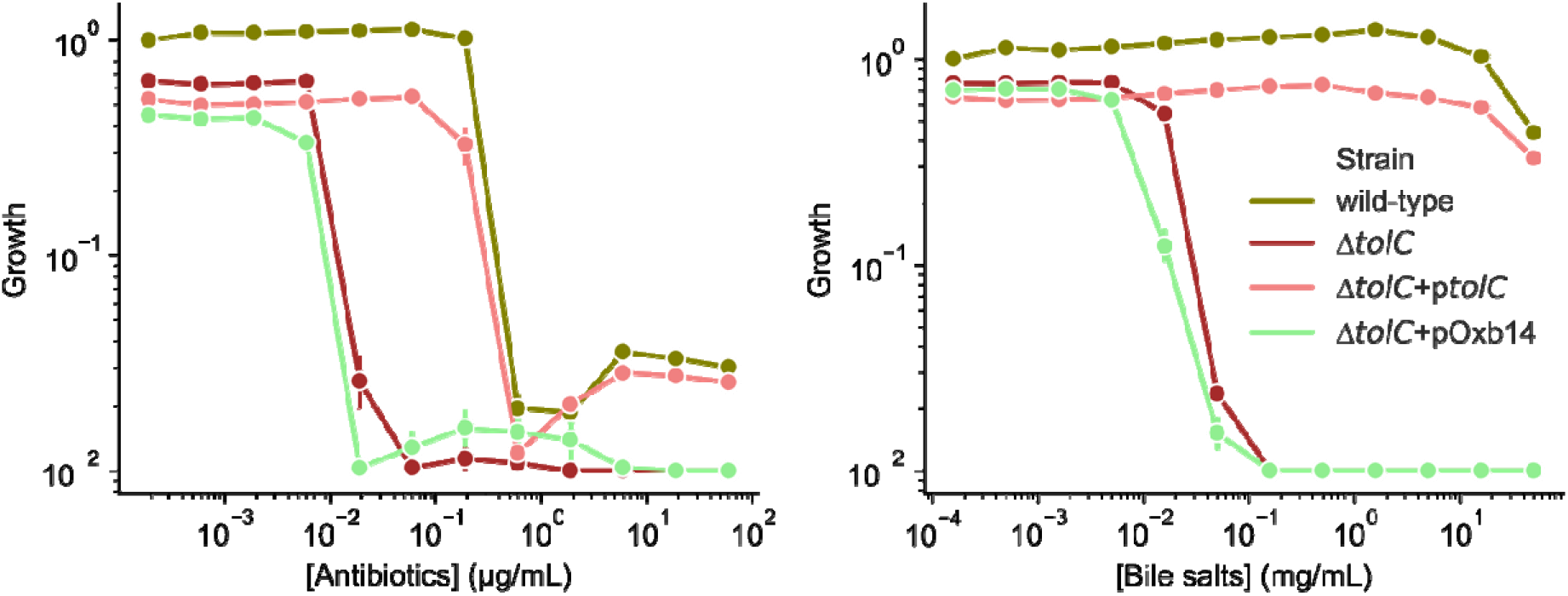
Supplementing the *tolC* gene into the *E. coli*-Δ*tolC* rescues antibiotic and bile salt resistance. Dose response curves are reported under antibiotics (left panel) and bile salts(right panel) selection for wild-type BW25113, ΔtolC, ΔtolC+pOxb14 and ΔtolC+ptolC strains. Growth of the strains are measured at 600nm every 30 minutes. Area under the growth curves are calculated for each antibiotic and bile salts concentration. Growth values are reported on y-axis after growth values are normalized using wild-type growth in the absence of selection.

**Figure S3.**
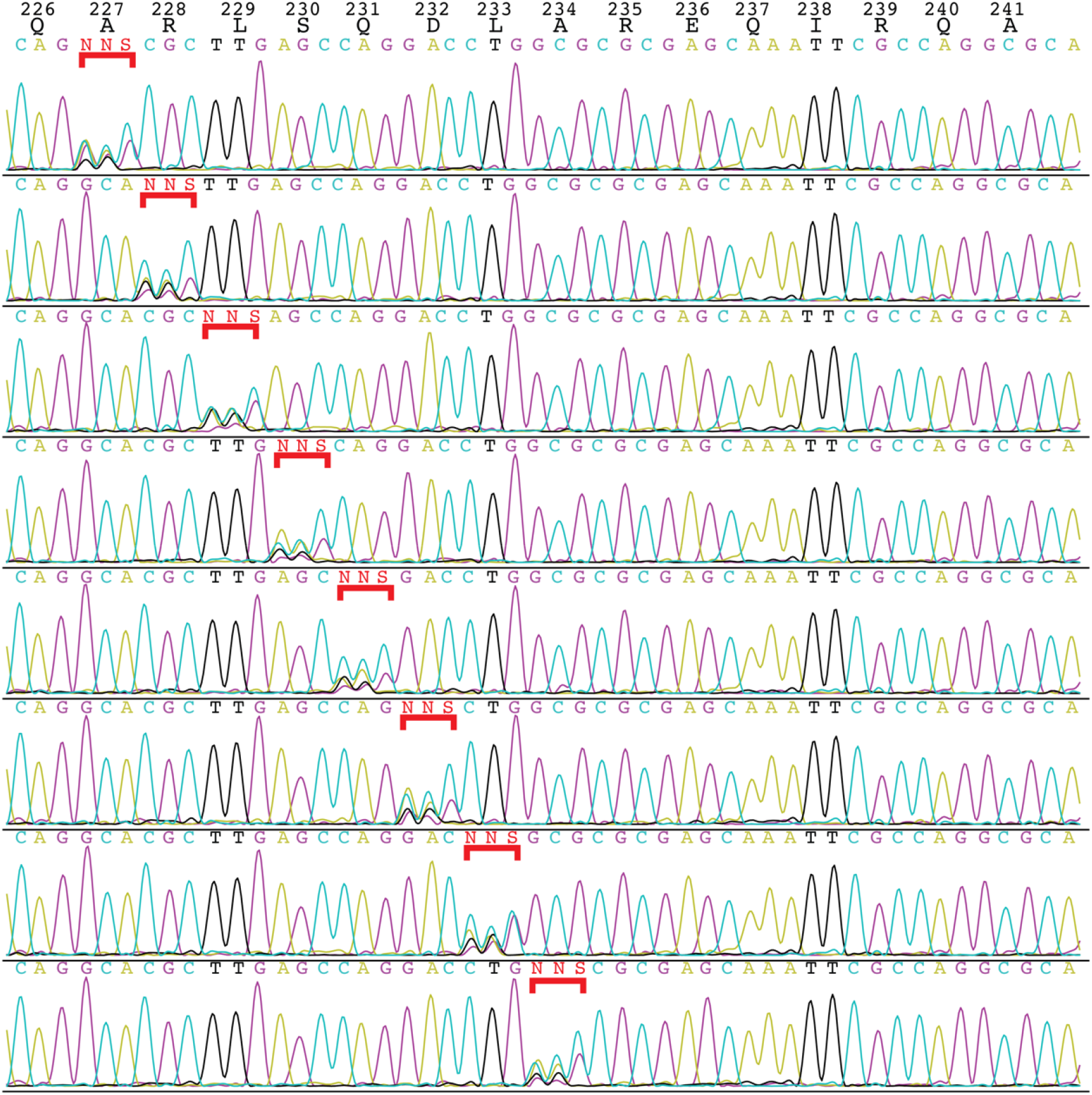
Confirmation of saturation gene mutagenesis with Sanger sequencing. Native amino acids for mutated positions (residues 227-234) are shown in black in the top row. Randomization of the intended residue was done separately for each position. Chromatograms demonstrate successful randomization of each codon to NNS (Any nucleotide in first two position and either G or C in the third position). In chromatograms, Cyan color was used for Cytosine, Black for Thymidine, Magenta for Guanine, and Yellow for Adenine. Sequence of each fragment is shown above the chromatogram. Randomized regions are highlighted using a red squared bracket.

**Figure S4.**
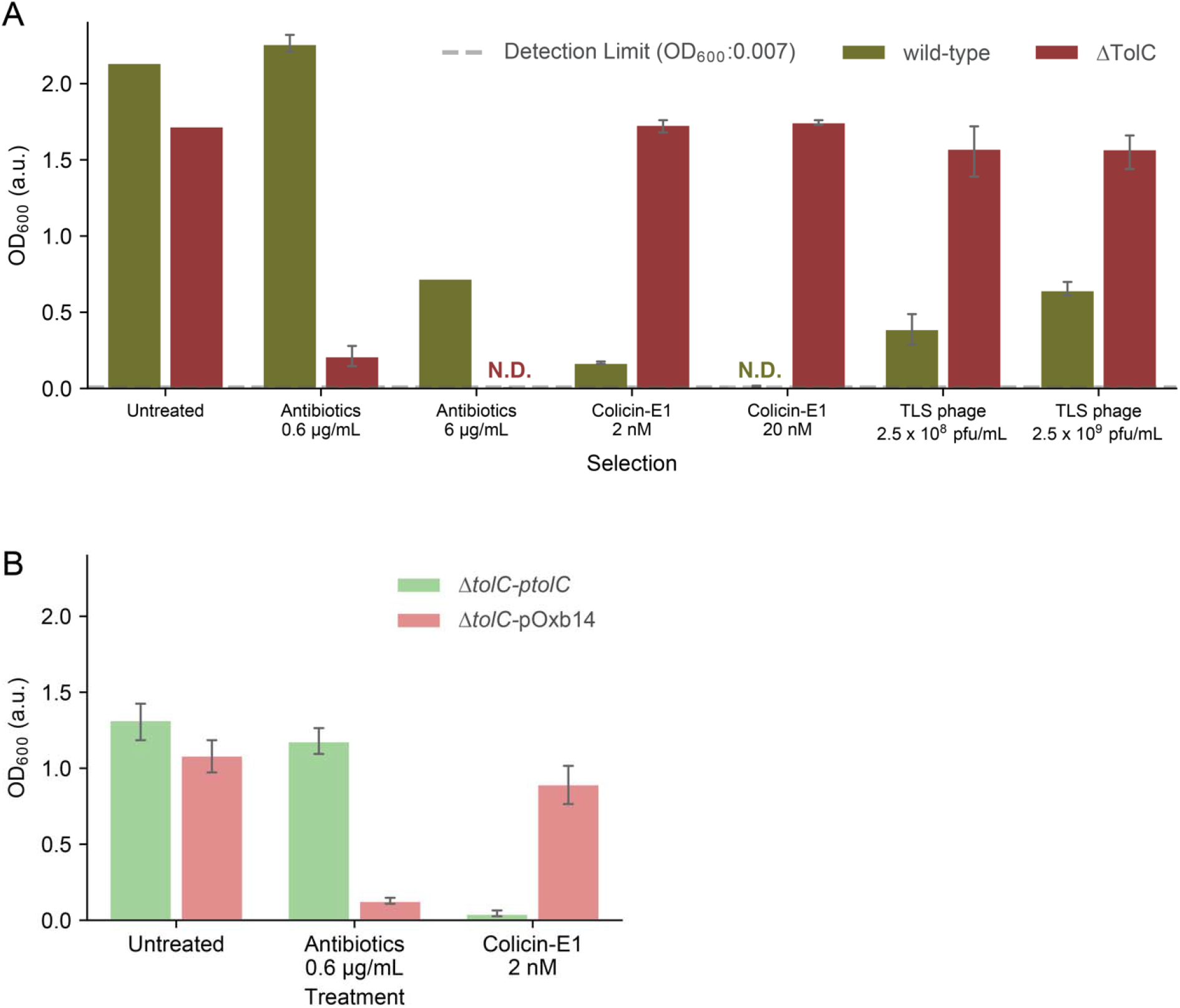
Effects of antibiotic, colicin-E1 and TLS phage selections are quantified with an optical density based assay for wild-type *E. coli* (green) and *E. coli*:Δ*tolC* (red). We plotted cell densities with and without selection. **(A)** Growth values at two concentrations of antibiotics, colicin-E1 and TLS phage were tested. No growth has been observed for ΔtolC strain in Antibiotics (6 μg/mL) selection and for wild-type strain for colicin-E1 (20nM) selection which are represented as not determined (N.D. with respective colors). Detection limit for spectrophotometer plotted as horizontal gray dotted line. **(B)** Growth is measured in the presence of antibiotics (piperacillin-tazobactam) and colicin-E1 for the *E. coli*:Δ*tolC* strain supplemented with an empty plasmid (red) and with a plasmid carrying the wild type *tolC* gene (green).

**Figure S5.**
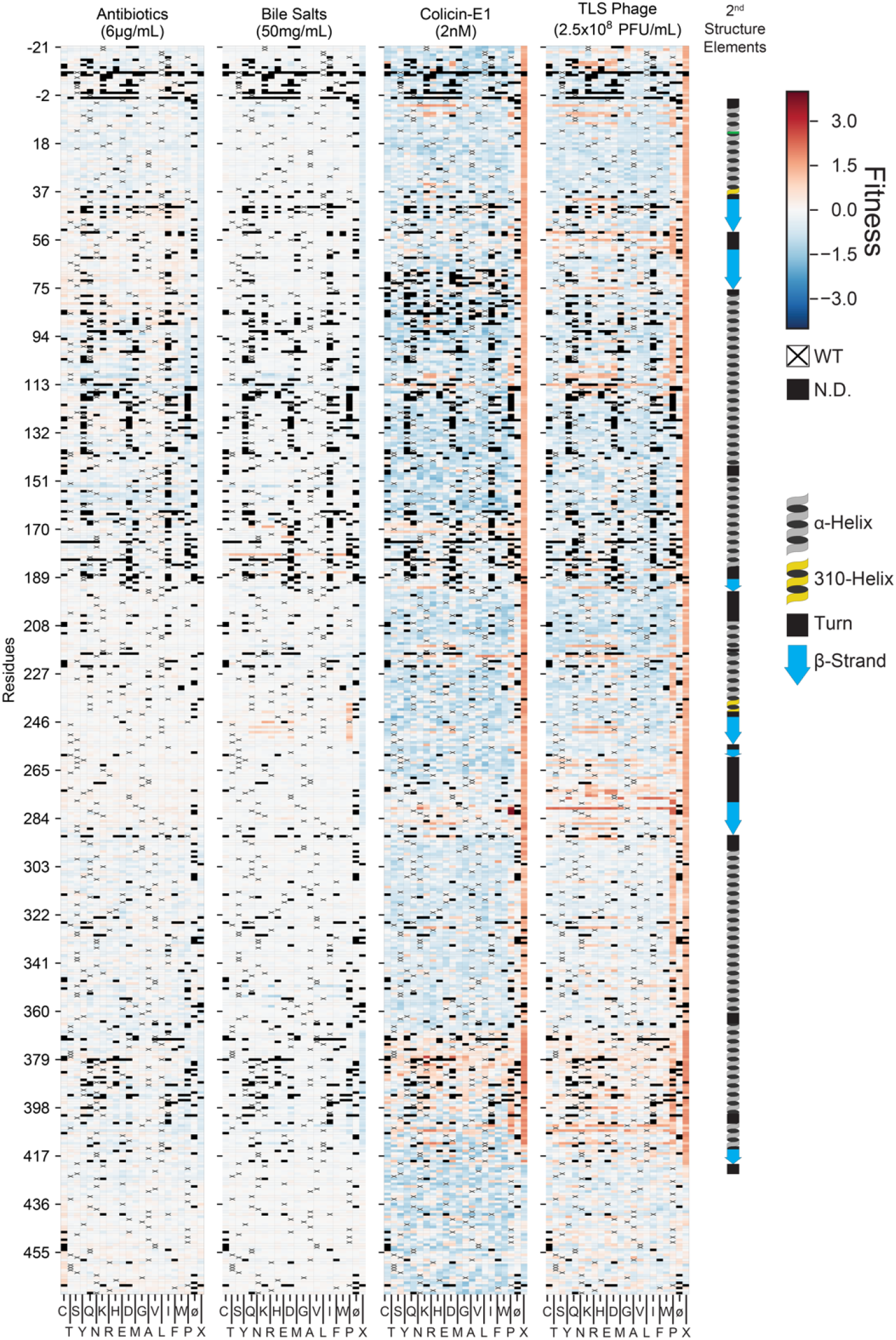
Fitness effects under four different stress conditions are plotted as heatmaps. Y axis in the heatmaps show the residues. Columns in heatmaps represents synonymous(∅), nonsynonymous and stop codon (X) mutations on each residue. Known secondary structure elements of TolC protein are highlighted in color.

**Figure S6:**
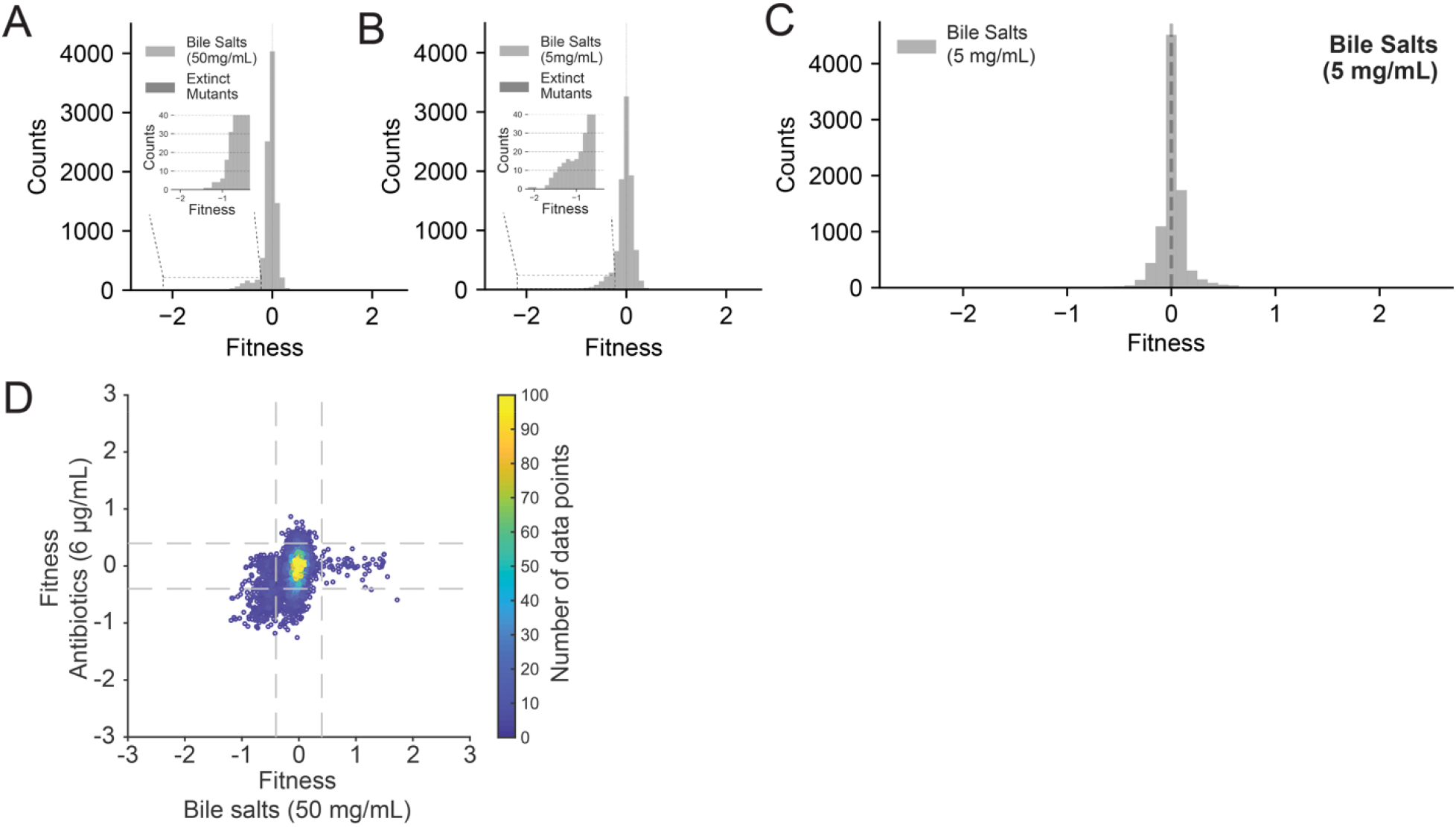
Distribution of fitness effects (DFEs) for bile salts selection. **(A-B)** DFEs were calculated under two different selection strengths. DFEs under bile salts selection are narrow and centered around neutrality (s = 0) regardless of the selection strength, with tails extending to the left (increased sensitivity, insets). (C) DFE of bile salts selection measured in Illumina NovaSeq platform. Distribution is narrow and centered around neutrality as well. (D) Comparison of fitness effects of TolC mutations under antibiotics (6μg/mL) and bile salts (50 mg/mL) selections. There is a weak correlation in fitness values under these selection conditions (*ρ*= 0.28 and p<0.001 Pearson Correlation). Vertical and horizontal dashed lines represent three standard deviation from mean (±0.4).

**Figure S7:**
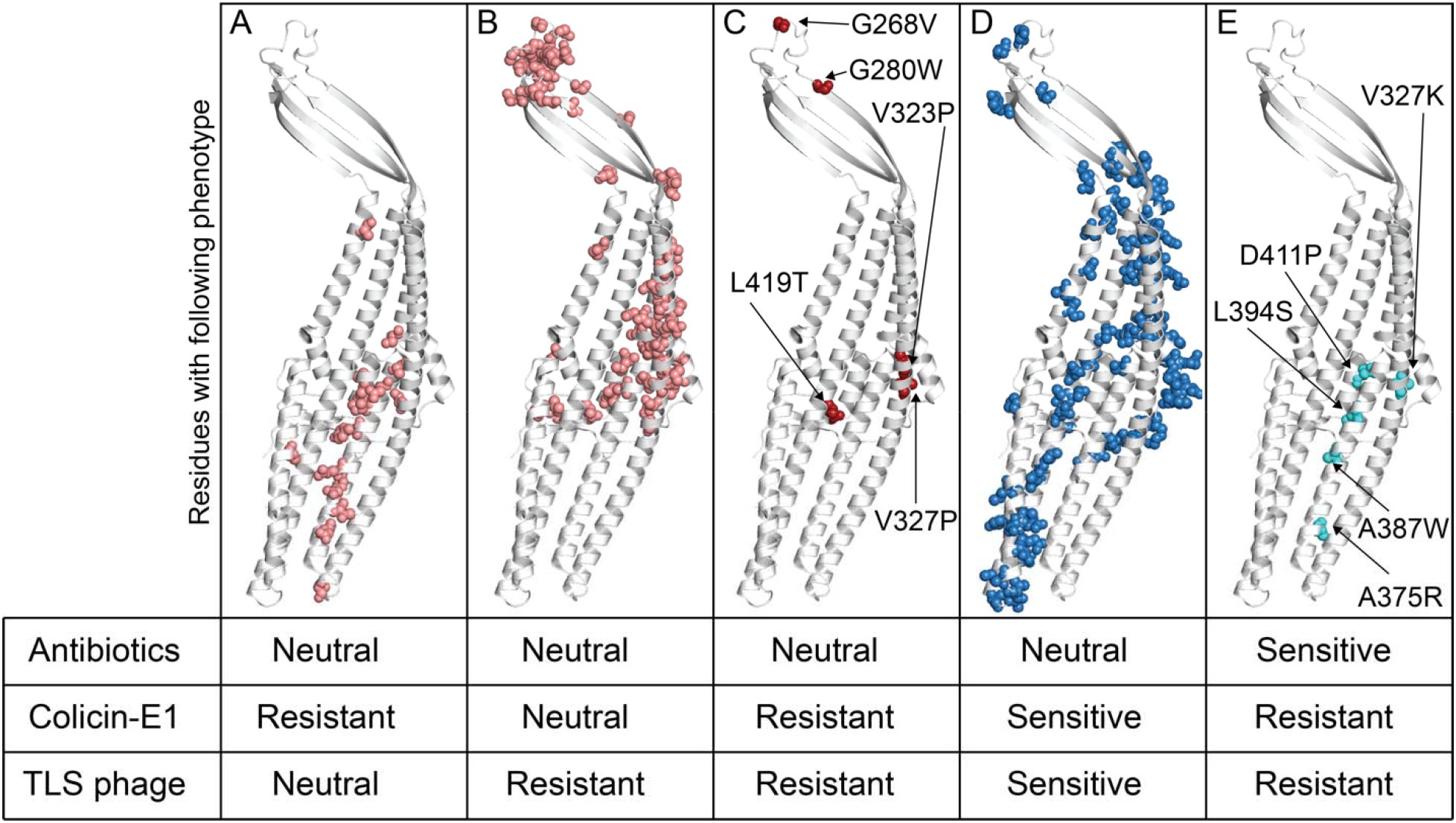
Mutated residues that caused significant fitness changes under at least one of the three selection factors are highlighted in color on monomeric TolC structure. (Table S2) **(A)** Mutations in pink colored residues confer resistance to colicin-E1 without causing significant fitness changes under other selection conditions. **(B)** Mutations in pink colored residues confer resistance to TLS phage without causing significant fitness changes under other selection conditions. **(C)** Mutations in red colored residues increase resistance to both colicin-E1 and TLS phage without disrupting efflux of antibiotics. **(D)** Mutations in blue colored residues increase sensitivity to both colicin-E1 and TLS phage without disrupting efflux of antibiotics. **(E)** Mutations in cyan colored residues increase resistance to both colicin-E1 and TLS phage and disrupt efflux of antibiotics. These mutations likely cause misfolding of TolC or blockage of the TolC channel as their fitness effects are reminiscent of the loss of the *tolC* gene.

**Table S1:** Mean and standard deviation values for DFEs under all selection conditions: antibiotics (0.6 and 6 μg/mL), bile salts (5 and 50 mg/mL), colicin-E1 (2 and 20nM), and TLS phage (2.5×10^8^ and 2.5×10^9^ pfu/mL).

**Table S2:** Fitness values of mutations represented in Figure S7 are tabulated. There are six sheets in the excel file. First sheet shows the consistent data that has similar fitness values in two different experiments. Remaining five sheets show fitness values for the mutations represented in Figure S7A-E.

